# Heterologous Expression and Characterization of *ToxA1* Haplotype from India and its Interaction with *Tsn1* for Spot Blotch Susceptibility in Spring Wheat

**DOI:** 10.1101/2023.04.17.536213

**Authors:** Ranjan Kumar Chaubey, Dharamsheela Thakur, Sudhir Navathe, Sandeep Sharma, Vinod Kumar Mishra, Pawan Kumar Singh, Ramesh Chand

## Abstract

**Background:** *ToxA*, a necrotrophic effector protein, is present in the genome of fungal species like *Parastagnospora nodorum*, *Pyrenophora tritici-repentis* and *Bipolaris sorokiniana. Tsn1* is the sensitivity gene in the host whose presence indicates more susceptibility to *ToxA* carrying pathogen, and *ToxA*-*Tsn1* interaction follows an inverse gene-for-gene relationship.

**Methods and Results:** The present study involved cloning and expressing the *ToxA1* haplotype from *B. sorokiniana*. It was found that the amplicon exhibited an expected product size of 471 bp. Sequence analysis of the *ToxA1* nucleotide sequence revealed the highest identity, 99.79%, with *P. tritici-repentis*. The protein expression analysis showed peak expression at 16.5kDa. Phylogenetic analysis of the *ToxA1* sequence from all the *Bipolaris* isolates formed an independent clade along with *P. tritici-repentis* and diverged from *P. nodorum*. *ToxA-Tsn1* interaction was studied in 18 wheat genotypes (11 *Tsn1* and 7 *tsn1*) at both seedling and adult stages, validating the inverse gene-for-gene relationship, as the toxin activity was highest in the K68 genotype (*Tsn1*) and lowest in WAMI280 (*tsn1*).

**Conclusion:** The study indicates that the haplotype *ToxA1* is prevailing in the Indian population of *B. sorokiniana*. It would be desirable for wheat breeders to select genotypes with *tsn1* locus for making wheat resistant to spot blotch.

## Introduction

Host selective toxins (HSTs) are the effector molecule(s) released during the invasion of necrotrophic pathogens and act as major determinants of pathogenicity (1). These necrotrophic effectors are specific biomolecules that are structurally complex, chemically diverse, and responsible for causing some key diseases in important crops. HSTs are very important in host-pathogen interaction but are produced by limited genera of fungi involving multiple toxins (2). Spot blotch of wheat is a damaging disease caused by *Bipolaris sorokiniana* (Sacc.) Shoemaker, syn. *Cochliobolus sativus* (S. Ito & Kurib.) a hemi-biotrophic pathogen. This disease is mainly prevalent in South Asia’s warm, humid region, especially in the eastern Indo-Gangetic plains of India. Wheat productivity can decline by 40% due to spot blotch alone (3). Spot blotch has been reported to cause up to 17.5% yield loss in the Indian sub-continent, and about 25mha of wheat is affected by spot blotch worldwide (4). Plant pathogens have their genetic makeup for causing host infection, which can decided by the presence of a single gene that confers host susceptibility or resistance (5, 6). The proteinaceous necrotrophic effector ToxA a major HST produced by *B. sorokininana*. Additionally, ToxA is known to be produced from two necrotrophic pathogens, *Parastagonospora nodorum* (Berk.) Quaedvl., Verkley & Crous, syn. *Phaeosphaeria nodorum* (E. Müll.) Hedjar. and *Pyrenophora tritici-repentis* (Died.) Drechsler and cause Septoria nodorum blotch and tan spot, respectively (7, 8). ToxA protein was reported initially in a culture filtrate of the wheat pathogen *P. tritici-repentis* (8). The ToxA protein was first discovered by Tomás and Bockus (9) from the culture filtrate of *P. tritici-repentis*. The *ToxA* manifested high nucleotide diversity in *P. nodorum,* whereas it showed a lack of variation in *P. tritici-repentis*. This pattern of nucleotide diversity and the recent emergence of tan spot suggests the recent introduction of *ToxA* into *P. tritici-repentis* genome (6). The *ToxA* sequences from *P. nodorum* and *P. tritici-repentis* depicted differences at four fixed nucleotide sites resulting in two predicted amino acid changes consistent with the interspecific transfer of this gene from *S. nodorum* into *P. tritici-repentis* (6), and this *ToxA* gene jumped horizontally from *P. tritici-repentis* to another pathogen *B. sorokiniana,* (10). *ToxA* and its surrounding 14 kb suggest that this gene was horizontally transferred, facilitated by a type II DNA transposon (11). *ToxA* is a single-copy gene, and it was found to be embedded within a 12-kb genomic element nearly identical to the corresponding regions in *P. nodorum* and *P. tritici-repentis* (10). The mature ToxA protein is 13.2 kDa, and it encodes a signal peptide of 23 amino acids and a pro-domain of 4.3 kDa (required for protein folding), and both are cleaved before the secretion of mature ToxA protein (1, 12, 13). Studies revealed that the host sensitivity of ToxA was mapped on wheat chromosome arm 5BL, and the sensitivity gene was named *Tsn1* (14). Further, it was confirmed that this gene was significant for both tan spot and SNB, and in both cases, the pathogen had a sensitivity gene such as *ToxA* (6, 15). For the wheat-*B. sorokiniana* pathosystem, ToxA confers virulence, and it is the only known virulence gene whose corresponding sensitivity gene (*Tsn1*) in the host has been identified. Further, it was also demonstrated that *Tsn1*-carrying wheat cultivars are more sensitive to *B. sorokiniana* isolates that harboured the *ToxA* gene. Cloning of the *Tsn1* gene unfolded characteristics similar to classical disease resistance nucleotide-binding leucine-rich repeat (NLR) gene that included serine/threonine kinase, leucine-rich repeats, nucleotide binding and protein kinase domains (14). The ToxA gene is transcribed as a necrotrophic effector protein responsible for infection in the host, with *Tsn1* being a resistance gene (16). A resistance-like gene *Tsn1* in wheat confers susceptibility to *ToxA* in *P. tritici-repentis* and *P. nodorum* isolates (6). This *ToxA–Tsn1* interaction system has been proved as an example of the inverse gene-for-gene relationship. It was realizing the occurrence of spot blotch in South Asia and the emergence of *ToxA1* as a key virulence gene driven us towards studying this host-pathogen relation. Keeping this in mind, we cloned the *ToxA1* gene from *B. sorokiniana* and conducted expression analysis for the ToxA1 protein. We also validated the pathogenicity of *ToxA1* on the allelic status of *Tsn1* by using 18 *Tsn1/tsn1* genotypes.

## Materials and Methods

### Primer designing

*ToxA* is a 534 bp-long gene sequence, and primers were designed using the web-based tool Primer-BLAST (Basic Local Alignment Search Tool) from NCBI. The *ToxA* sequence from *P. tritici-repentis* (U79662.1) was obtained in FASTA format from NCBI (National Centre for Biotechnology Information) GenBank database (https://www.ncbi.nlm.nih.gov/genbank/). The primers were utilized to amplify *ToxA* in *B. sorokiniana* isolate MCC 1533 (Pusa2). The isolate MCC1533 (Pusa2) has been identified and deposited at National Center for Microbial Resources (NCMR), Pune, India. The primer sequence used is given as Forward Primer (FP) - CGCGGATCCGATCCCGGTTACGAAATCG and Reverse Primer (RP) - CGC CTCGAGCTAATTTTCTAGCTGCATTCTC. Revised nomenclature of ToxA haplotypes corresponds the

### RNA extraction and cDNA synthesis

*The B. sorokiniana* isolate MCC 1533 (Pusa2) was cultured for 10 days in a Potato Dextrose Broth. Mycelium was harvested, and the total RNA was isolated using Invitrogen PureLink^TM^ RNA Mini kit following the manufacturer’s protocol. The integrity and size distribution of total RNA purified with RNeasy Kit was checked by 1.2% agarose gel electrophoresis and ethidium bromide staining, visualizing two sharp bands of 28S rRNA to 18S RNA. RNA was quantified using Nanodrop2000 instrument (Thermo Scientific) and 1µg RNA reverse transcribed to cDNA was synthesized using a Thermo cDNA synthesis kit following the manufacturer’s protocol.

An additional treatment of RNase was included during cDNA synthesis as supplied and instructed with the kit and manufacturer’s protocol to prevent chances of contamination of genomic DNA during isolation.

### ToxA gene cloning

The expression construct was produced in the bacterial expression vector pET28a that directs the expression of HIS-tagged fusion proteins in *E. coli*. BamH1 and XhoI restriction sites were introduced in forward and reverse primer for cloning in pET vector. Primers were designed such that the PCR amplified portion of the *ToxA1* gene was generated with a 5’ BamHI site and a 3’ XhoI site. The PCR product was digested with BamHI and XhoI, gel-purified with the QIAquick Gel Extraction Kit (Qiagen, Valencia, CA) and cloned into the BamHI and XhoI sites of pET28a. Primers FP-CGCGGATCCGATCCCGGTTACGAAATCG and RP-CGC CTCGAGCTAATTTTCTAGCTGCATTCTC were used for the PCR amplification. PCR amplification was performed using cDNA as a template in an Eppendorf Mastercycler using the following cycles; initial denaturation of 95°C for 7 min followed by 35 cycles of 94°C (30 s), 55°C (45 s) and 72°C (1 min) and a final extension of 10 min at 72°C. The PCR product was gel-purified, digested with BamHI and XhoI, and ligated into the respective restriction sites of pET28a. The resulting construct contained the predicted effector protein gene fused to a 6X histidine tag at the N-terminus. The ligated product was used to transform *E. coli* DH5alpha and BL21 (DE3) strains (17) for cloning and protein expression, respectively.

### Sequencing and characterization

Cloned fragment of *ToxA* from *B. sorokiniana* isolate MCC1533 (Pusa2) was sequenced by Sanger sequencing. The sequences obtained in two frames were merged, and Bioedit Sequence Alignment Editor (v 7.2) was used to identify overlapping sequences in both orientations (18). Integrated sequences were then prepared in a single frame. The sequence has been deposited to the National Centre for Biotechnology Information (NCBI) database with Gene Bank accession number (OP289649). The processed *ToxA* sequence was used as a query for BLASTn (https://blast.ncbi.nlm.nih.gov/Blast.cgi) for finding homologous sequences with reference to the NCBI nucleotide database (19). The processed *ToxA1* nucleotide sequence was then translated to the amino acid sequence in BioEdit 7.2. Sequences of *P. tritici-repentis, P. nodorum* and different isolates of *B. sorokiniana* were retrieved from the NCBI protein database. The amino acid sequences retrieved using BLASTp searches were then used to create multiple sequence alignments of 27 ToxA sequences. MEGA X (Molecular Evolution Genetics Analysis) software (19) was then used to generate a phylogeny of *ToxA* amino acid sequences of 49 isolates of *P. tritici-repentis, P. nodorum* and *B. sorokiniana* using the CLUSTAL W alignment algorithm (20). All alignment parameters were left at their default values; 178 and 114 positions with and without gaps were obtained, respectively. Sequences were analyzed using substitutions selected with complete deletion of gaps or absent data. The phylogenetic tree was inferred using the JTT model and the maximum likelihood method (21). Using the default setting and the Neighbor Joining method, the initial tree was inferred, and Nearest-Neighbor Interchange was used as the ML heuristic search method. Phylogenetic tree reconstructions were performed using the Maximum Likelihood Method. For the Poisson correction model, 1000 bootstrap replicates were used to assess node support (22).

### Protein modelling and topographical properties

The predicted protein sequence of cloned Pusa2 *ToxA* was subjected to the SWISS-MODEL homology modelling online tool (23). Proteins were modelled utilizing Protein Data Bank (PDB) entry 1zld.1.A (*P. tritici-repentis*) as reference. The quality of modelled protein was judged based on GMQE (Global Model Quality Estimation) and QMEAN (Qualitative Model Energy Analysis) Z-score parameters (24, 25). The best model was selected based on the GMQE score, which ranges from 0 to 1, with a higher value indicating better reliability. On a global scale, the QMEAN Z-score estimates the “degree of nativeness” of the structural features observed in the model. QMEAN Z-scores around zero indicate good agreement between the model structure and experimental structures. Scores of -4.0 or below are an indication of models with low quality. The quality of the resulting models was also monitored with PROCHECK and ProsSA-WEB (26). Geometrical and topographical properties of the modelled proteins were identified using the Computed Atlas of Surface Topography of Proteins (CASTp) 3.0 online server (27). The UCSF Chimera software was used to create images of molecular graphics (28). The alpha helix and the beta-sheet percentage were analyzed using PDBMD2CD.

### Isolation of proteins from E. coli culture

*E. coli* colony harboring the expression construct was inoculated in 5 ml of LB (Sambrook et al. 2006) with 100μg of kanamycin per ml (LBA) and grown overnight at 37°C and shaking at 250 rpm (G24 Incubator Shaker, New Brunswick Scientific, Edison, NJ). The overnight culture was used to inoculate 25 ml of fresh LBA, which was then incubated with shaking at 37°C for one hour. The expression of fusion proteins was induced by the addition of isopropyl-beta-Dthiogalactopyranoside (IPTG) to a final concentration of 1 mM, followed by an additional 4 hours of incubation under the same conditions. After 20 minutes of centrifugation at 4,000 g and 4°C, cell particles were resuspended in a cold sonication buffer(SB; 50 mM Na2HPO4, 5 mM Tris-base, 300 mM NaCl, pH 8.0) with 0.5% Triton-x-100 and 10 mM DTT. Resuspended pellets were sonicated three times, 30 s each, with 2 min on ice between periods. Insoluble material was then pelleted by centrifugation at 10,000 × *g,* 4°C, for 10 min. Pellets were washed thrice with SB with 0.5% Triton-× -100 and 10 mM DTT. The final washed pellets, containing inclusion bodies, were either stored frozen at –80°C or resuspended in 1 ml of 8 M guanidine HCl (Sigma) in refolding buffer (250 mM NaCl, 100 mM sodium phosphate buffer, 10 mM Tris-base, 4% glycerol, 1 mM EDTA, and 0.005 % Tween 20, pH 8.0) with freshly prepared DTT added to a final concentration of 10 mM. Solubilized proteins were sonicated briefly twice (for 5 to 10 s) and then heated to 37°C for 10 min and 50°C for an additional 5 min to assist in the solubilization of proteins. Insoluble material was removed from the samples by centrifuging at 12,000 ×*g* for 10 min. Then, solubilized proteins were quantitated with Bradford assay reagent (Bio-Rad, California, USA) with bovine serum albumin used as a standard. Fusion proteins solubilized in guanidine HCl were used immediately.

### Screening of ToxA1 –Tsn1 interaction on different wheat genotypes

The screening of 18 wheat genotypes was conducted for *Tsn1/tsn1,* out of which 11 genotypes harboring *Tsn1* and the rest 7 genotypes with *tsn1* (see Table 1) were sown in 3 replications at polyhouse, Plant Pathology, Institute of Agricultural Sciences, Banaras Hindu University in 1m rows at a distance of 22.5cm between two rows. Infiltration of ToxA1 protein, as well as control (Uninduced protein), was done at both the seedling stage (3-5 leaf stage) and adult stage (Zadoks growth stage 55) (29). 250µg of ToxA protein was infiltrated by a sterilized syringe, and symptoms were recorded 7 days after infiltration. The 5^th^ emerged leaf was infiltrated for seedlings, while the flag leaf was infiltrated with toxin ToxA1 for adult plants. The temperature was maintained at 25±2℃. The light/dark period was maintained at 14hrs light and 10hrs of darkness. Photographic data was captured using a Nikon camera. The lesion size was measured in length, width, and area of lesions using ImageJ software (Version 1.53s) (30).

**Table 1:** Response of wheat genotypes (*Tsn1/tsn1*) in terms of leaf necrosis after infiltration of ToxA1 protein at both seedling and adult stage of wheat genotypes harboring Tsn1/tsn1 allele.

| Genotypes | Length(cm) |  | Width(cm) |  | Area(cm2) |  | Tsn1/tsn1 |
| --- | --- | --- | --- | --- | --- | --- | --- |
|  | Seedling | Adult | Seedling | Adult | Seedling | Adult |  |
| **K 68** | 2.872 | 4.613 | 0.404 | 0.530 | 0.977 | 2.326 | *Tsn1* |
| **C 306** | 1.921 | 4.226 | 0.329 | 0.633 | 0.678 | 1.737 | *Tsn1* |
| **HUW468** | 2.057 | 3.472 | 0.402 | 0.438 | 0.718 | 1.407 | *Tsn1* |
| **Yangmai 6** | 0.860 | 1.406 | 0.311 | 0.381 | 0.316 | 0.450 | *Tsn1* |
| **K65** | 1.513 | 2.582 | 0.380 | 0.566 | 0.388 | 1.096 | *Tsn1* |
| **WAMI160** | 1.615 | 3.461 | 0.345 | 0.506 | 0.646 | 1.521 | *Tsn1* |
| **WAMI188** | 1.693 | 2.582 | 0.310 | 0.500 | 0.585 | 1.036 | *Tsn1* |
| **WAMI224** | 1.643 | 2.857 | 0.330 | 0.488 | 0.476 | 1.298 | *Tsn1* |
| **WAMI226** | 1.782 | 2.320 | 0.405 | 0.503 | 0.628 | 1.090 | *Tsn1* |
| **WAMI277** | 1.464 | 2.174 | 0.360 | 0.485 | 0.464 | 1.024 | *Tsn1* |
| **WAMI288** | 1.591 | 2.054 | 0.432 | 0.575 | 0.622 | 1.017 | *Tsn1* |
| **Max.** | 2.872 | 4.613 | 0.432 | 0.633 | 0.977 | 2.326 |  |
| **Min.** | 0.860 | 1.406 | 0.310 | 0.381 | 0.316 | 0.450 |  |
| **HUW234** | 0.964 | 1.894 | 0.335 | 0.408 | 0.345 | 0.561 | *tsn1* |
| **PBW343** | 0.860 | 1.446 | 0.349 | 0.408 | 0.319 | 0.423 | *tsn1* |
| **LOK1** | 1.098 | 1.761 | 0.307 | 0.432 | 0.278 | 0.493 | *tsn1* |
| **Sonalika** | 1.919 | 2.923 | 0.361 | 0.492 | 0.584 | 1.321 | *tsn1* |
| **WAMI280** | 0.972 | 1.305 | 0.319 | 0.327 | 0.265 | 0.411 | *tsn1* |
| **WAMI289** | 1.065 | 1.471 | 0.319 | 0.344 | 0.342 | 0.529 | *tsn1* |
| **WAMI290** | 1.206 | 1.467 | 0.327 | 0.360 | 0.365 | 0.493 | *tsn1* |
| **Max.** | 1.919 | 2.923 | 0.361 | 0.492 | 0.584 | 1.321 |  |
| **Min.** | 0.860 | 1.305 | 0.307 | 0.327 | 0.265 | 0.411 |  |

### Statistical Analysis

The statistical analysis was conducted for phenotypic traits, including length, width, and area of necrotic lesions of wheat genotypes with the *Tsn/tsn1* gene at both seedling and adult stages. The point biserial correction coefficients between disease components and the presence/absence of *Tsn1* were calculated (32). The analysis in R software V 4.2.1 was performed using packages’ *phytools’, ’stats’, ’agricolae’, ’ggplot2’, ’ggtree’* (31).

## Results

### Cloning and expression study of ToxA1

The *ToxA* gene from *B. sorokiniana* was amplified using primers designed from *PtrToxA* through RT-PCR. The amplicon exhibited the expected product size of 471bp, which was then cloned in the pET28a vector. The cloned gene was validated using colony PCR of the transformed cell. The *ToxA1* gene, after validation, was subjected to sequencing, and the sequence data after processing revealed successful cloning of the *ToxA* gene (now *ToxA1*-*B. sorokininana*). This *ToxA1* nucleotide sequence was further used for BLASTn analysis, revealing the highest identity, 99.79%, with *P. tritici-repentis* and E-value close to zero. Expression of ToxA1 was done by inducing the transformed *E.coli* cell through IPTG of different concentration, namely, at 0.4mM, 0.6mM, 0.8mM and 1.0mM. Maximum expression of ToxA1 was obtained at 1.0mM concentration with an expected band size of 16.5kDa in sodium dodecyl sulfate (SDS)-polyacrylamide gels (Fig. 1).

**Figure 1.**
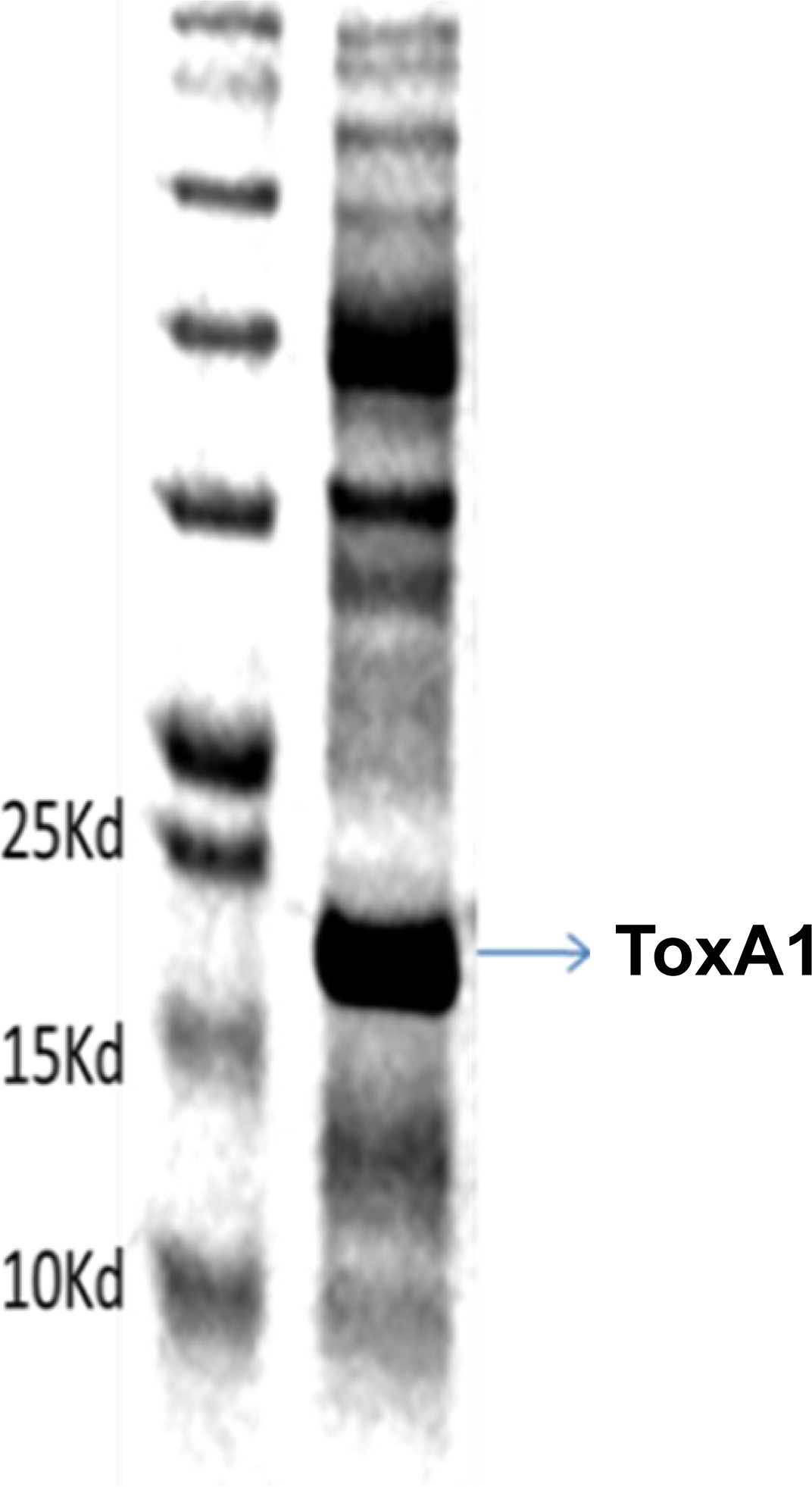
Expressed ToxA1 protein showing 16.5kd band size on SDS-PAGE

### ToxA1 characterization and Phylogenetic analysis

The identification of *ToxA1* after showing sequence similarity with *P. tritici-repentis ToxA1* using BLASTn analysis on NCBI was conducted. Further characterization of ToxA1 was conducted by translating the nucleotide sequence to amino acid sequence using the web-based tool Expasy (https://www.expasy.org/resources/translate). The amino acid sequence of ToxA1 was subjected to BLASTp analysis on NCBI, which depicted 100% similarity with *P. tritici-repentis* showing E value 9e^-109^. Multiple sequence alignment of 25 ToxA protein sequences and *ToxA1* MCC1533 (Pusa2) translated sequence in Bioedit revealed that amino acid differences were found in most *P. nodorum* strains. However, the cloned Pusa2_*ToxA1* sequence showed complete similarity to *P. tritici-repentis* ToxA sequence. *P. nodorum ToxA* sequence depicted amino acid substitution for Glutamate, Aspartate, Leucine, Valine and Asparagine (Fig. 2).

**Figure 2.**
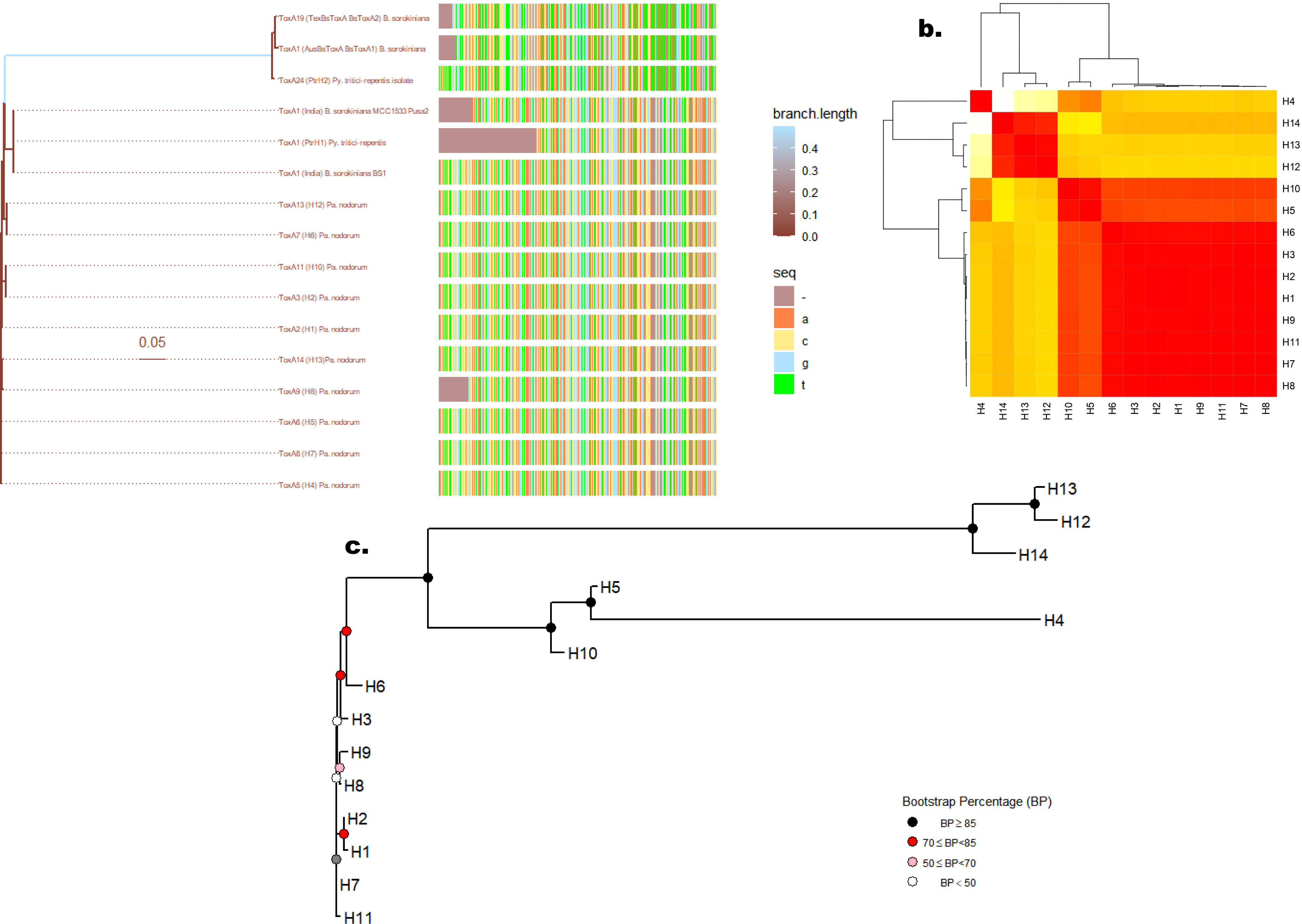
(a.) The multiple sequence alignments with a phylogenetic tree, Colors both on the phylogenetic tree and the branch scale represent genetic distance. ToxA sequences have been coloured by four colours shown in the ’seq’ column for each nucleotide. Colour changes on the aligned sequences represent nucleotide differences. (b) Based on the number of nucleotide variations between ToxA haplotypes, a heat map was created. The relevant ToxA haplotype in the matrix is represented by each branch of the phylogenetic tree. The near ties to “dark red” and the distance ties to “white” colour. (c) Neighbor-joining (NJ) tree for ToxA haplotypes based on the nucleotide sequence differences. Coloured internal nodes represent the bootstrap confidence level.

The phylogenetic analysis depicts the evolutionary relatedness of the *ToxA1* amino acid sequence with other ToxA sequences. The tree indicates that *ToxA1* sequences originate from a common ancestor but form different clades during evolution. The *ToxA1* sequence from all *Bipolaris* isolates formed an independent clade along with *P. tritici-repentis* and diverged from *P. nodorum*. The phylogeny indicates sequence similarity among *B. sorokiniana* and *P. tritici-repentis,* whereas it differed from *P. nodorum* (Fig. 3).

**Figure 3.**
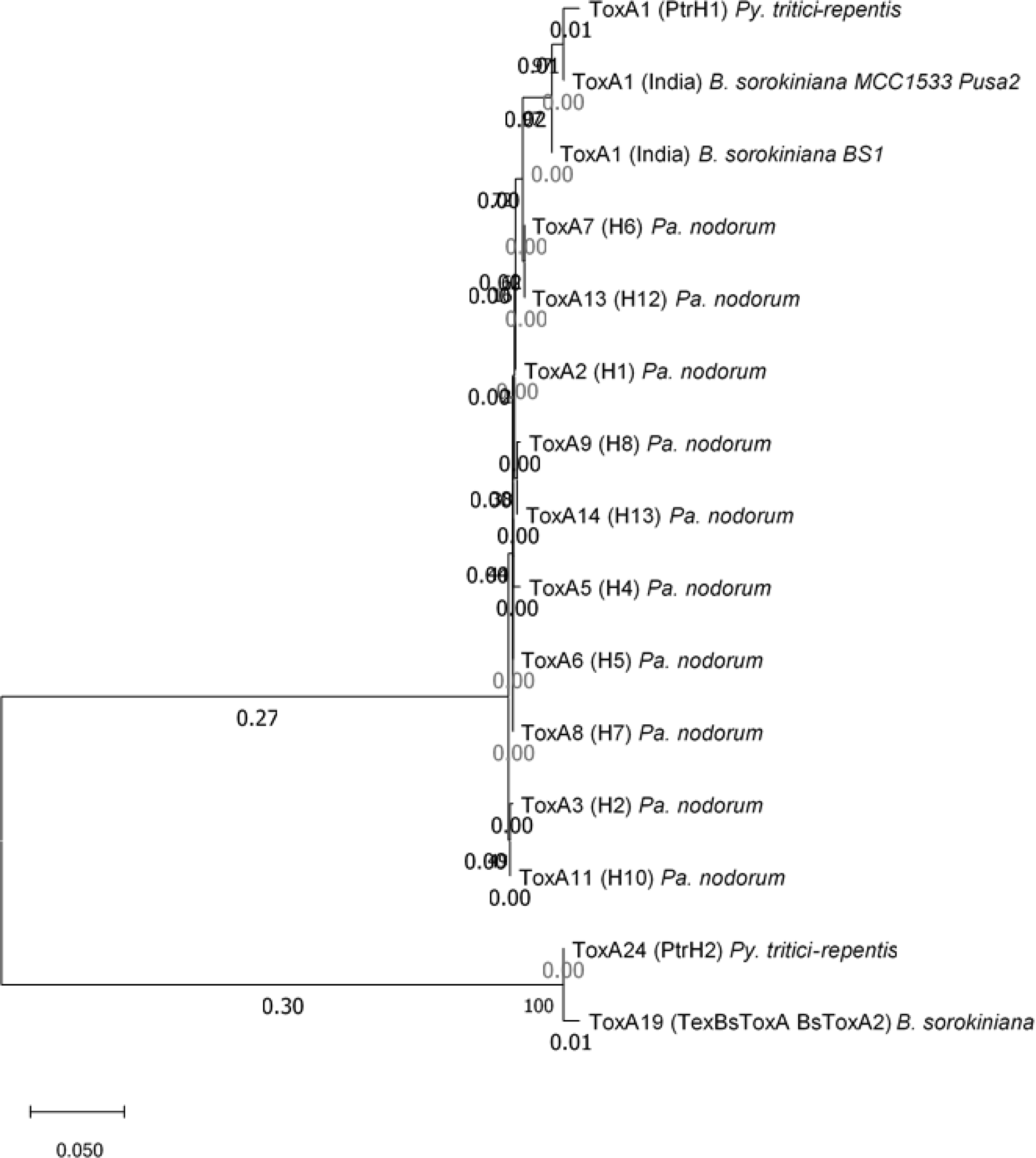
Phylogenetic tree using Maximum Likelihood algorithm of different ToxA1 of *Pyrenophora tritici-repentis, Bipolaris sorokiniana,* and varipus haplotypes of *Parastagnospora nodorum*. The percentage of bootstrap support (1000 replicates) is indicated at corresponding nodes. The nomenclature of the haplotypes is based on the Aboukhaddour et al. (2023)

### Protein modelling and Topographical properties

The structure of the ToxA1 protein of a cloned gene of *B. sorokiniana* isolate MCC1533 (Pusa2) was predicted using the SWISS-MODEL homology modelling online server. For checking the reliability of models GMQE (ToxA1: 0.56), QMEAN Z scores are included QMEAN -0.33, Cbeta –1.79, torsion 0.53 and these scores indicated that these models are reliable and good quality. The quality of the resulting models was monitored with PROCHECK. Ramachandran plot analysis revealed that 88.1% of the ToxA1 protein structure residues fell within the most favoured regions, with a further 11.9% in the additionally allowed region (Fig. 4). No residues were found in the generously allowed or disallowed regions. The number of non-glycine and non-proline residues was 100%. The number of end residues (excluding Gly and Pro) is 2, and the number of glycine and proline residues was 12 and 3, respectively.

**Figure 4.**
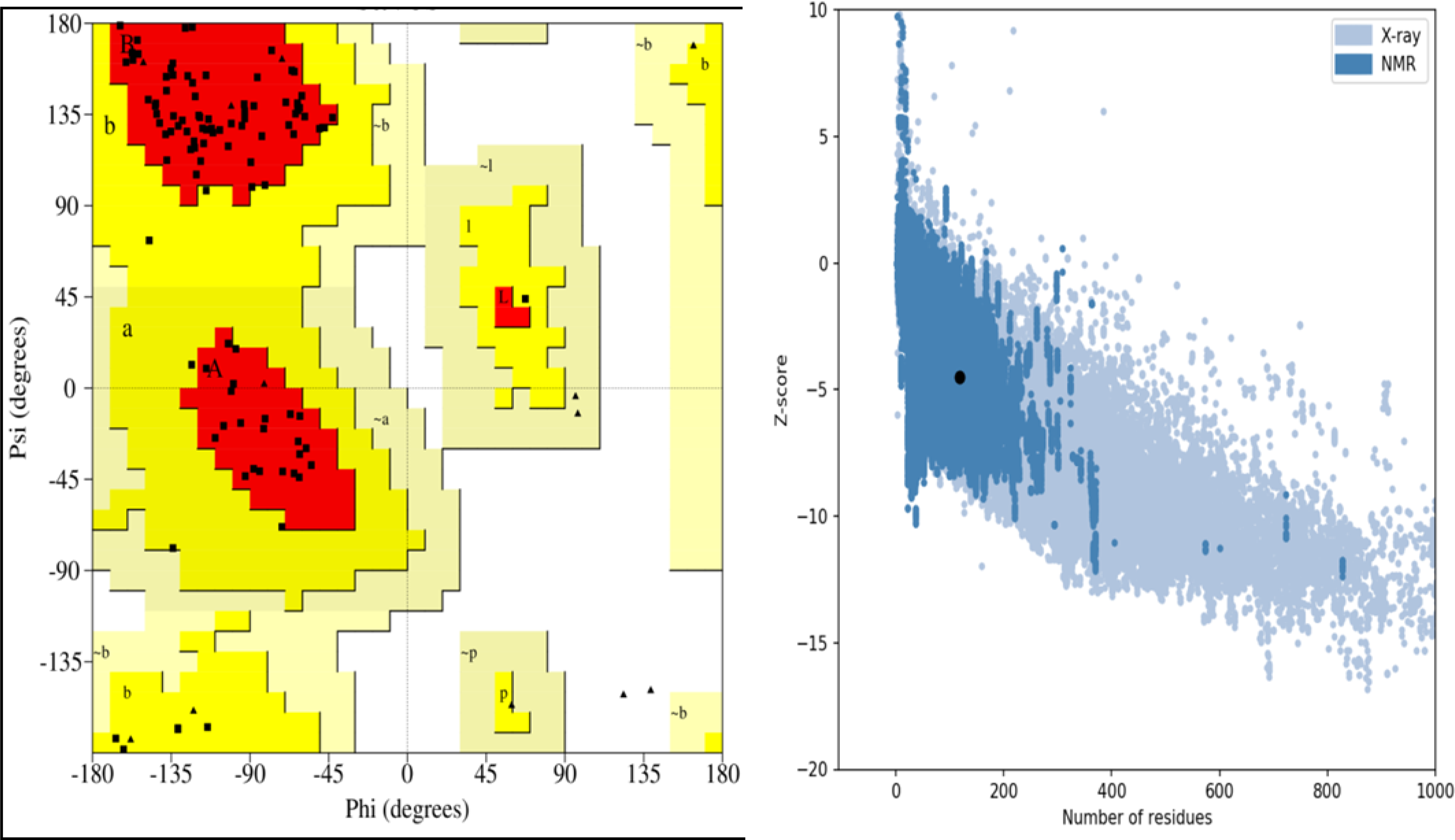
Ramachandran plot and ProsSA z-score analysis of modelled ToxA1 protein from *B. sorokiniana*. Ramachandran distributions of amino acids in the model were calculated using pro check (a), and the z-score plot was plotted using ProsSA-WEB (b).

Predicted ToxA1 protein structures of *B. sorokiniana* MCC1533 (Pusa2) isolate based on homology modelling are depicted in Fig. 4. The protein sequences of ToxA1 protein showed sequence identity of 100% with template PDB entry 1zld.1.A (*P. tritici-repentis*, *PtrToxA*). When generated structures were compared in UCSF Chimera, it was observed that the RGD motif (toxic region) was present in the 3D structure of ToxA1 protein and found that arginine is present in the alpha helix of the loop at 118 positions. At the same time, Glycine and Aspartic acid is present in the β-sheet of the protein structure at 119 and 120 positions of ToxA1 protein (Fig. 5). The percentage of the alpha helix was 3.39%. At the same time, antiparallel sheet 1 was 35.55%, antiparallel sheet 2 was 12.76%, the parallel sheet was 0.85%, the turn was 7.63%, and the other was 39.83%.

**Figure 5.**
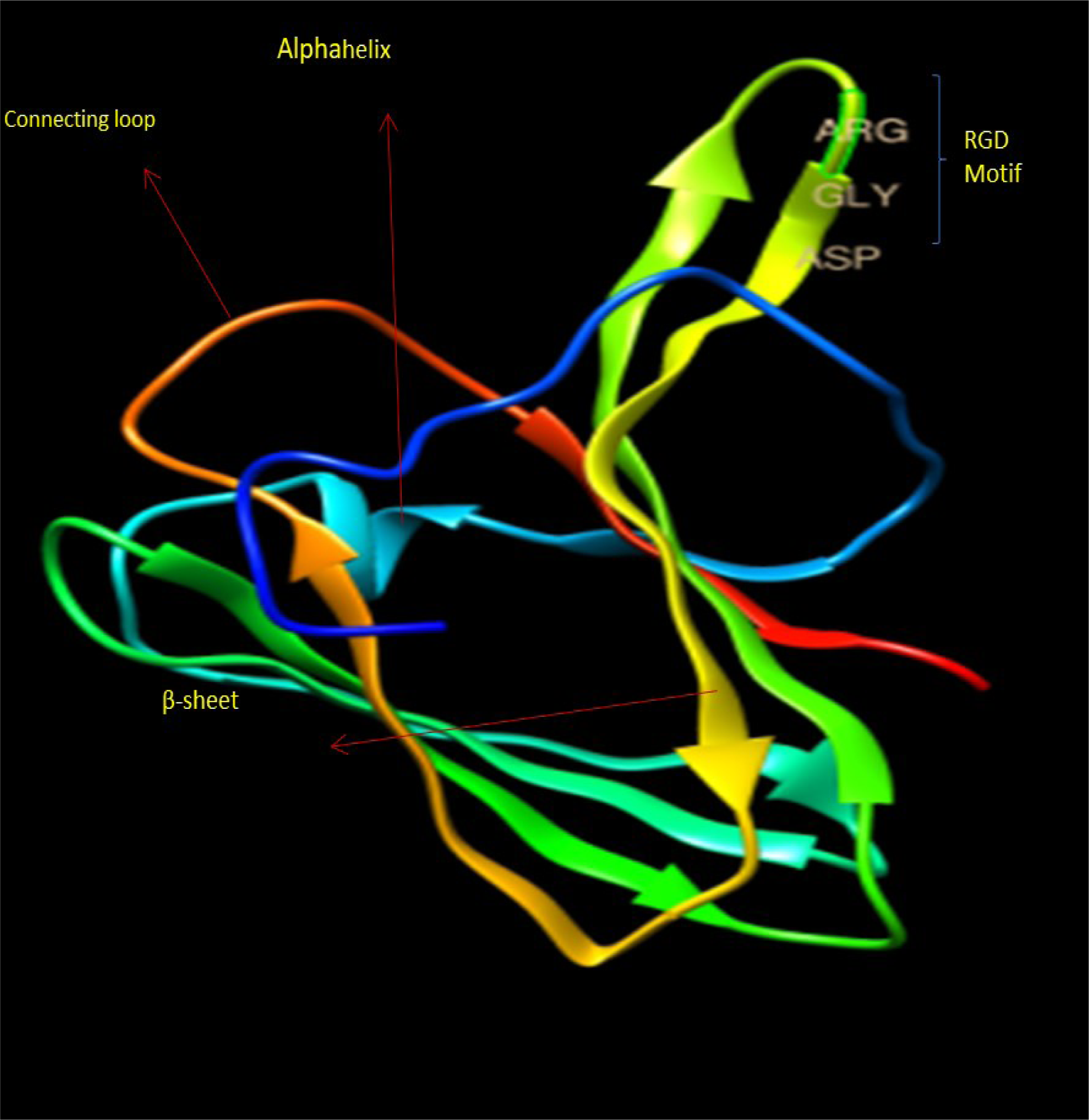
*ToxA1* protein structure of *B. sorokiniana* isolate MCC 1533 (Pusa2) depicting conserved RGD motif (ARG-GLY-ASP)

**Figure 6.**
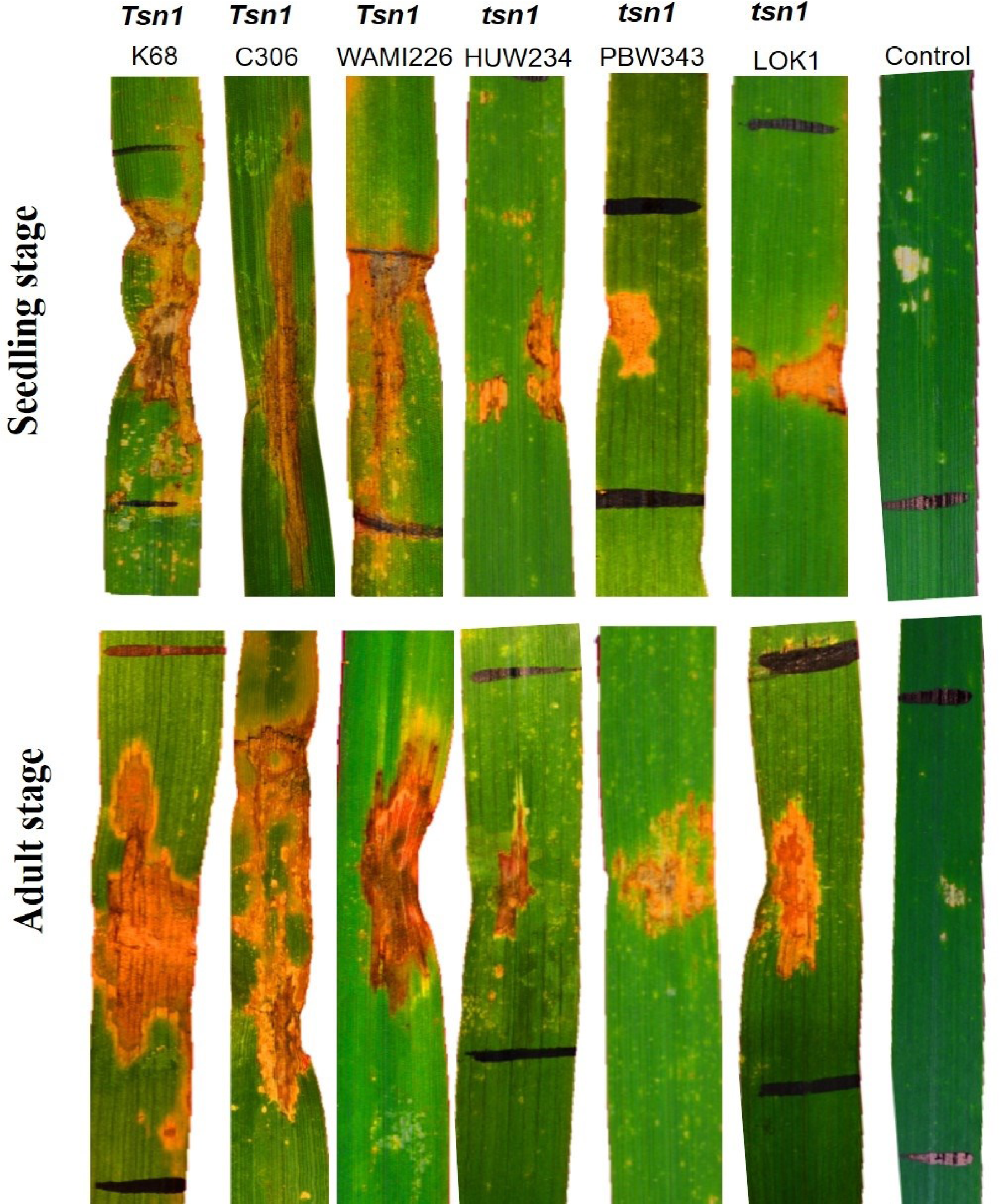
Symptom of leaf necrosis on *Tsn1/tsn1* harbouring genotypes after infiltration ToxA1 protein at seedling (3-5 leaf stage) and on flag leaf of adult plant (Zadoks growth stage 55).

### Bioactivity of ToxA1 effector protein and its interaction with Tsn1 at both seedling and adult stage

The bioactivity of the *ToxA1* effector was studied on necrotic lesions symptoms caused due to *ToxA1* protein and was found to be more prominent at the adult stage. At the seedling stage, the maximum symptomatic area (0.977 cm^2^) was recorded in genotype K68(*Tsn1*) and the minimum (0.265 cm^2^) in WAMI280 (*tsn1*). At the same time, the length varied from 2.872 cm (K68, *Tsn1*) to 0.860cm (PBW343, *tsn1*). However, the maximum width recorded was 0.432 cm for genotype WAMI288 (*Tsn1*), and the minimum in genotype harbouring *tsn1*(LOK1) was 0.307cm.

At the adult stage, the length of the symptom was maximum (4.613cm) in genotypes harbouring*Tsn1*(K68) and minimum (1.305 cm) WAMI280 (*tsn1*), and the width also had the same trend ranging from 0.633cm (C306, *Tsn1*) to 0.327cm (WAMI280, *tsn1*). The area was measured for quantifying the necrotrophic symptom, and it found that the maximum area in the genotype harboring *Tsn1* was 2.326 cm^2^ (K68), and the minimum was 0.411 cm^2^(WAMI280, *tsn1*). A significant difference was found between the seedling and adult stages and for *Tsn1* and *tsn1* (Table 1, Fig. S2, S3, S4).

## Discussion

Studies on necrotrophic effectors producing HSTs have provided a new dimension to host-pathogen interaction and established an inverse gene-for-gene relationship (6, 34). The HSTs are effectors proteins and induce a susceptible response in the host harbouring the dominant susceptibility gene. ToxA is a necrotrophic effector protein, one of the key factors for causing the most devastating disease, spot blotch in wheat, prevalent in warm, humid regions of South Asia. The present study focuses on cloning of *ToxA1* gene, followed by its characterization and purification of the resultant protein and its expression analysis. This study successfully cloned and characterized the *ToxA1* gene from *B. sorokiniana* isolate MCC1533 (Pusa2). This gene was found initially in *P. nodorum* and horizontally transferred to *P. tritici-repentis* and, more recently, to *B. sorokiniana* (6, 35). The cloned gene and its expression was studied in the present study, and the purified protein showed peak expression at a concentration of 1.0 mM IPTG with band migration at 16.5kDa in size. Manning et al. (36) also reported similar findings where lysates with His-ToxA1 expressed a phosphorylated band that migrated at 16 kDa compared to lysates without His-ToxA1 (control). However, mass spectral analyses have indicated a 13.2 kDa mass of purified mature ToxA1 protein from *P. tritici-repentis* (1, 8). The importance of a protein purification study for ToxA1, which has selective host toxicity for genetic analysis of pathogenicity in the *Pyrenophora*-wheat interaction, was also suggested by Touri et al. (8).

The *ToxA1* gene was truncated in the present study to remove the signalling sequences comprising 22 amino acids, as the mature ToxA1 gene is released after the proteolytic cleavage of this signal sequence. Previous studies have also described these signalling peptides in ToxA protein that involve the first 22 amino acids, which are proteolytically cleaved after import into the endomembrane system (1). The release of mature toxin involves additional proteolytic processing, and as the ToxA1 protein is blocked at the N terminus, such modifications would incur a mass increase (1). The ToxA1 cloned sequence obtained in our study has been compared with the already reported ToxA1 sequences from *P. nodorum*, *P. tritici-repentis* and *B. sorokiniana*. Multiple sequence alignment was performed on the amino acid sequences of 26 isolates. The results indicate that the ToxA regions of *B. sorokiniana* and *P. tritici-repentis* are more closely related to one another than they are to *P. nodorum*. This conclusion is supported by the identical ToxA1 sequence shared by *B. sorokiniana* and *P. tritici-repentis* and the six-position difference between *P. nodorum* haplotypes. (Fig. 2). The sequence identity between the three fungal species accounts for the common evolutionary origin of *ToxA* and the potential of exchanging DNA between these species (10). The significance of phylogenetic studies in establishing evolutionary relationships among the different pathogen races or deriving the specific gene evolution between different genera is well known. The sequencing data for *ToxA1* showed a high similarity between *B. sorokiniana* and *P. tritici-repentis* (10, 37, 38). Previous studies have also suggested that the ToxA1 sequence did not differ for *B. sorokiniana* and *P. tritici-repentis* but differed from the ToxA sequence of *P. nodorum* (10, 34, 37, 38). This evolutionary relatedness of *P. tritici-repentis* and *B. sorokiniana* sequences have also been reported earlier (10, 34). The earliest report suggested that ToxA has been present in *P. nodorum* for a long time as it exhibited greater sequence diversity and was interspecifically transferred to *P. tritici-repentis* quite later. This is evident from monomorphism or lesser sequence diversity in the ToxA1 sequence from *P. tritici-repentis* and the emergence of tan spot symptoms more recently in the near past (38).

Further insights into the ToxA structural analysis were performed by predicting the 3D protein structure using an amino acid sequence of ToxA1 (Fig.4). This is the first report for the 3D protein structure ToxA1 haplotype from *B. sorokiniana*. The predicted protein structure was similar to a single-domain protein with a β-sandwich fold of PtrToxA. This β-fold represents a solvent-exposed loop containing arginyl-glycol-aspartic acid (RGD)-functionally active motif (39, 40). ToxA1 has been known to resemble with fibronectin III domain which retains a solvent-exposed loop containing RGD motif for cell wall-plasma membrane interaction in plants (40, 41). It has been proposed that ToxA with RGD-containing motif has been known to interact with wheat mesophyll cells and thus establish disease (42). The mesophyll cells have a high-affinity matrix to recognize these motifs, and these all suggest the importance of the RGD motif in receptor binding and internalization into wheat cells (40).

ToxA is the only known virulence gene for which a host sensitivity gene (*Tsn1*) has been known in wheat-*B. sorokiniana* pathosystem. For the first time, the present study reports a cloned and purified protein ToxA1 infiltration in both seedling and adult stages of wheat lines harboring *Tsn1/tsn1*. The severity was enhanced in the adult stage compared to the seedling stage in *Tsn1* wheat lines. The present data also reveals that *Tsn1*-carrying genotypes exhibited more disease in seedlings than in the adult stage. The toxin activity was highest in the K68 genotype (*Tsn1*) and lowest in WAMI280 (*tsn1*) at both the seedling and adult stages. This could be validated by past studies suggesting that ToxA causes severe necrotic lesions on *Tsn1,* harboring wheat genotypes.

In contrast, less or no lesion was observed for wheat genotypes lacking the sensitivity gene (43). Navathe et al. (34) also reported one such possibility of the involvement of a recessive allele of *Sb2* towards an unknown HST. *Tsn1* has been reported attributing susceptibility towards both *P. nodorum*-causing SNB (6, 44) and *P. tritici-repentis*, causing tan spots (45). However, the environmental requirements for *Bipolaris* are different from these two pathogens, which are not prevalent in the South-Asian region.

## Conclusions

Nevertheless, it would be desirable for wheat breeders to select *tsn1* for making crops resistant to *B. sorokiniana, P. tritici repentis* and *P. nodorum* in nearest future. The study in this horizon should focus on loci other than *Tsn1* and *Sb2*. It would be interesting to study other quantitative trait loci involved to understand the complexity of this *ToxA-Tsn1* interaction and spot blotch disease resistance.

## Acknowledgements

Authors acknowledge Banaras Hindu University and Bihar Agricultural University for the infrastructure facility

## Author Contributions

Conceptualization, RC, VKM, PKS, SN, and SK; methodology, RKC and DT; validation, DT and SS; formal analysis, DT and SS; investigation, RKC; resources, RC, DT; data curation, DT, SS; writing—original draft preparation, RKC, SN; writing—review and editing, RC, PKS; supervision, RC, VKM, SS; project administration, RC; funding acquisition, RC, PKS. All authors have read and approved the manuscript.

Availability of data and materials: Not applicable.

## Declarations

### Ethics approval and consent to participate

Not applicable.

### Consent for publication

Not applicable.

## Supplementary Materials

**Figure S1.**
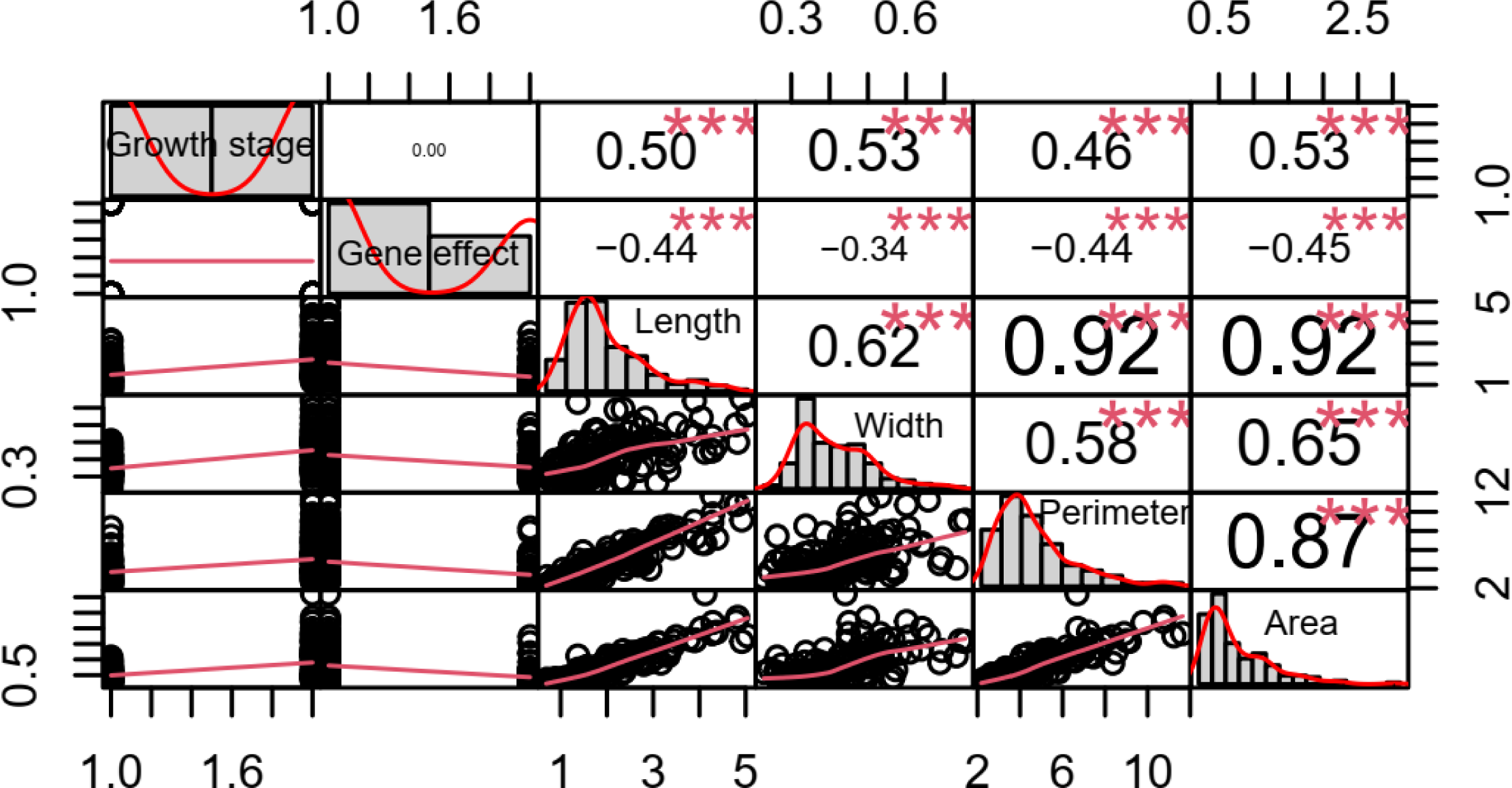
Correlation coefficients estimated by point biserial correlation for phenotypic traits associated with*Tsn1/tsn1* gene effect in wheat genotypes with ToxA1 isolate (MCC1533/Pusa2).

**Figure S2.**
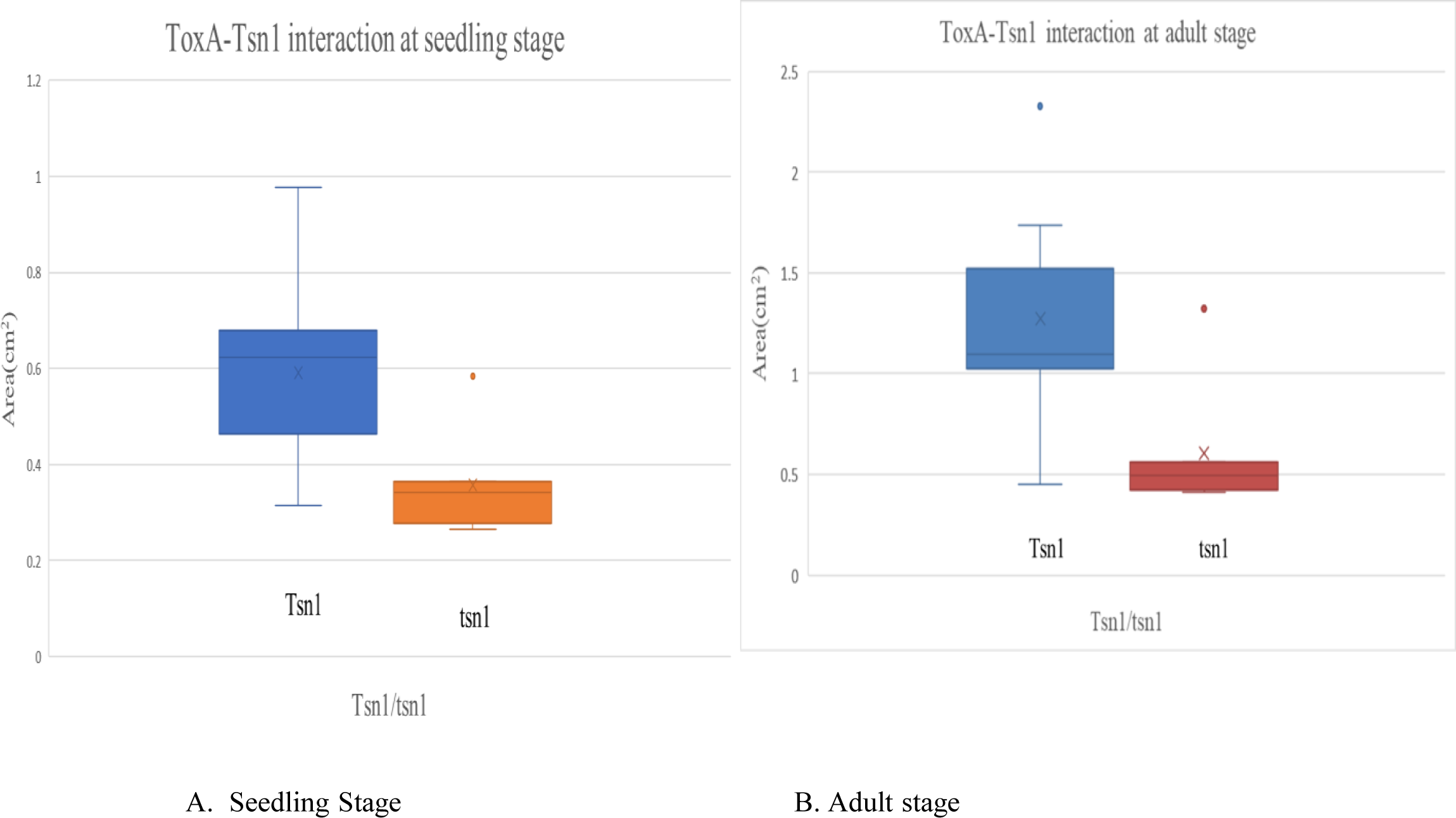
The necrotrophic area developed due to ToxA1–Tsn1 interaction at seedling (3-5 leaf stage) and adult stage (Zadoks growth stage 55) of 11 wheat genotypes with *Tsn1* and 7 wheat genotypes with *tsn1*.

**Figure S3.**
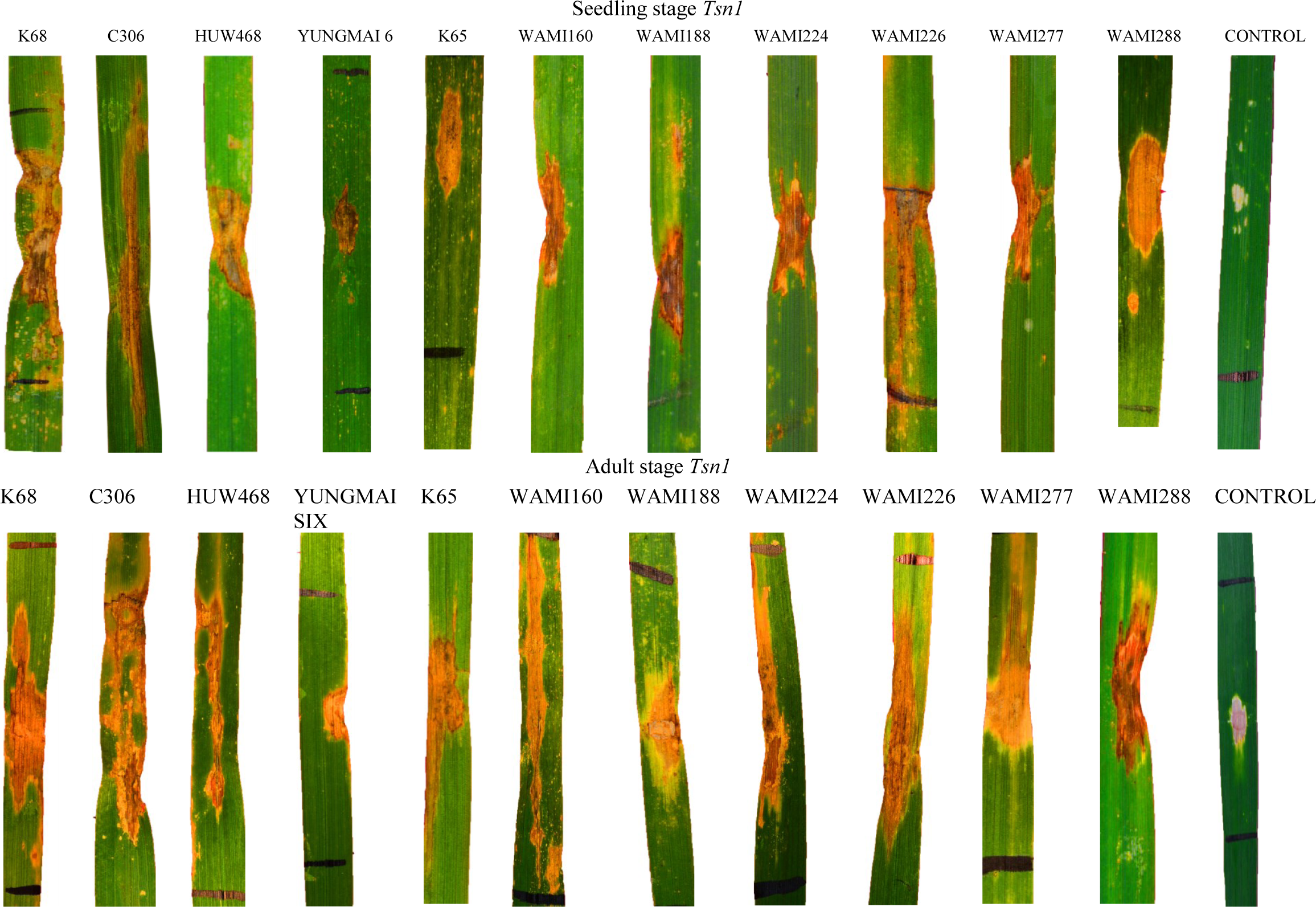
Symptom of leaf necrosis after infiltration ToxA1 protein at both seedling (3-5 leaf stage) and adult stage (Zadoks growth stage 55) of wheat genotypes harbouring *Tsn1*.

**Figure S4.**
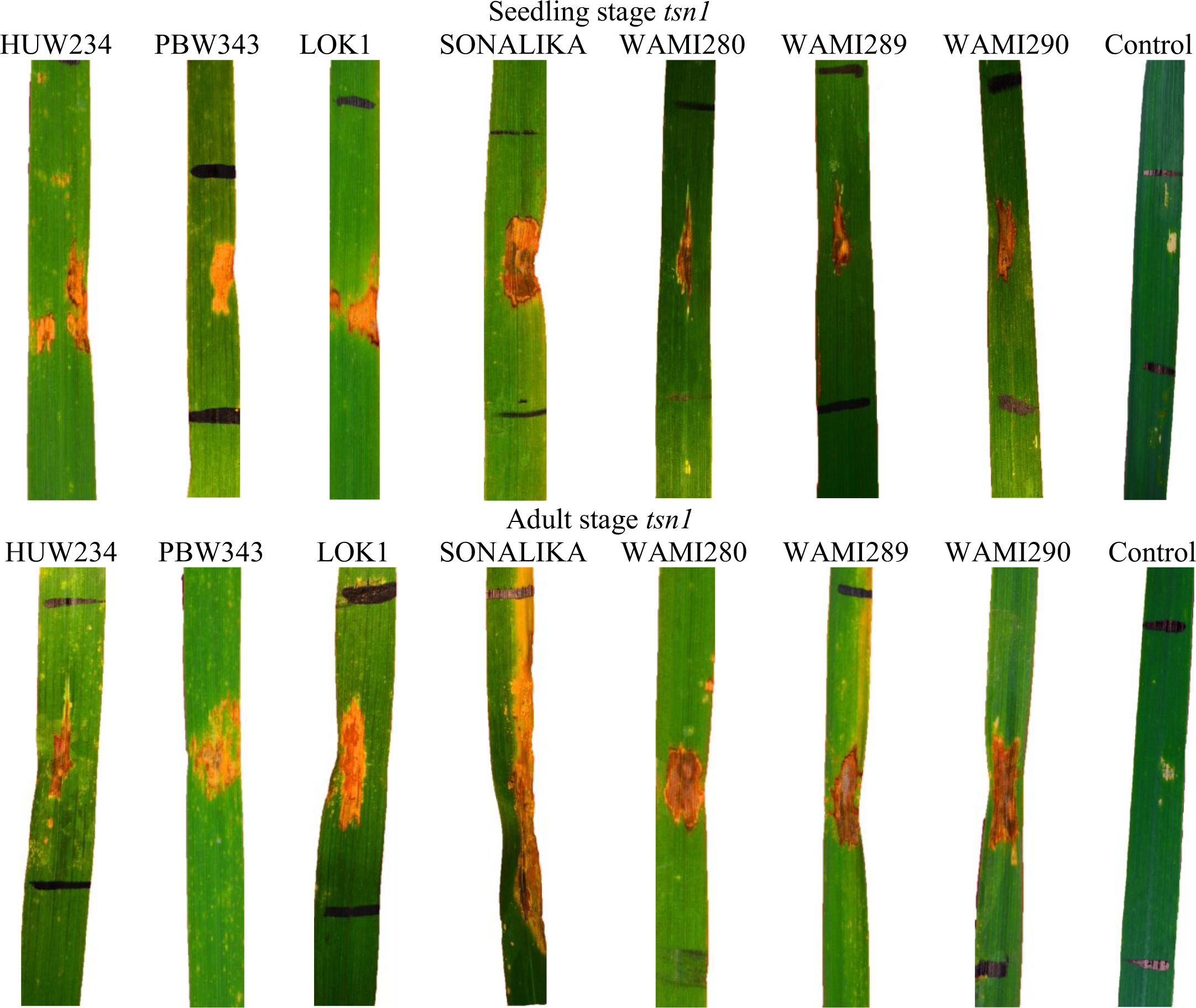
Symptom of leaf necrosis after infiltration ToxA1 protein at both seedling (3-5 leaf stage) and adult stage (Zadoks growth stage 55) of wheat genotypes harbouring *tsn1*.

